# Climate change and the thermal performance of a high protein food source: *Wolffia globosa*

**DOI:** 10.1101/2025.03.04.641473

**Authors:** K. Cuddington, M. Kuntze, D. Andrade-Pereira, Y. Gasmen, J. Wu, A. Ferns, X. Geng

## Abstract

*Wolffia globosa* is a tropical duckweed native to Southeast Asia, where it is harvested for food. This aquatic plant has a fast growth rate, a high protein content, and is also a source of important nutrients. Therefore, it could play an important role in food security under climate change and population growth. We provide the first thermal performance curve for *W. globosa,* and use this data to understand how climate change impacts on temperature in Southeast Asia may affect production. We find that the maximum relative growth rate occurs at constant temperatures of ∼32C. We find no significant difference between growth at current mean conditions and temperatures predicted in the next 40 years according to the high emissions scenario (SSP5-8.5 scenario) in Thailand, Laos and Myanmar when temperatures are held constant. However, the thermal performance curve is best described as asymmetric, with growth rates that fall rapidly at temperatures above this optimum. As a result, when temperatures are allowed to fluctuate about the mean in a pattern similar to recent heatwave conditions in Thailand, we find significantly lower growth rates at the optimum than at current mean temperatures. This decrease is driven by a significant increase in frond death at higher temperatures. However, given the fast growth rate of this species relative to other food crops, and the mitigating impact of water on the magnitude of temperature fluctuations, it seems likely that *W. globosa* may more rapidly recover from extreme heat events than other crop species. Therefore, it is likely to be a suitable candidate for adapting to climate change impacts.

## Introduction

In many regions, climate change is expected to increase both the mean and variance of air temperatures, simultaneously generating a higher frequency of heatwaves. Predictions regarding the potential impacts of these changes on food crops can be generated using thermal response studies that measure aspects of performance, such as relative growth rates or yield, as a function of temperature. The probable negative impacts on many crops and concurrent population growth both point to the need to identify species that may help with future food security. We examine the effects of temperature on the growth rates of *Wolffia globosa*, an aquatic plant that is a good source of protein and other nutrients. This data on thermal performance is used to determine optimal growing temperatures and to examine the probable impacts of changes in mean temperature and heatwaves on cultivation in southeast Asia.

*W. globosa* is a small, fast-growing aquatic plant that primarily reproduces asexually. Like other duckweed species, individual plants are recognized by their simplified, oval-shaped stem known as a frond that floats on the surface of calm freshwater in ponds, lakes, and marshes (Vu et al., 2020) *W. globosa* is native to Southeast Asia (Landolt 1986). However, it is also found in California, Africa and has more recently been reported in parts of Europe, including the Ukraine, the Czech republic (Vávra et al. 2024), the UK (Lansdown et al. 2022), Germany (Frank et al. 2020) and France (Lecron et al. 2021). *W. globosa* has traditionally been used as a food crop in some regions of Southeast Asia, such as Laos and northern Thailand (Baek et al. 2021), where it is sold under the local names khai nam, kai-pum, or kai nhae (literally meaning: water-eggs). The species is primarily harvested, rather than cultivated, twice a week from November to July (European Food Safety Authority 2021). (Note: According to Appenroth et al. (2017), the plant that has been widely consumed as food in Southeast Asia is *W. globosa* rather than *W. arrhiza* s mentioned by Bhanthumnavin and McGarry (1971)).

*W. globosa* has a very fast growth rate and a high protein content, which suggests that it could play an important role in sustainable food security (e.g., Nirmal et al. 2024). Boonarsa et al. (2024) report that the amount of protein in *W. globosa* harvested from farms in Northern Thailand is significantly higher than that in wheat, corn, brown rice and peas. In addition, the species has an abundance of other beneficial nutrients, and, unlike other plant foods, can provide vitamin B12, probably through endophytic bacteria (Kaplan et al. 2019). There is a proprietary strain of *W. globosa*, patented in 2018 (https://patents.google.com/patent/USPP29977P3/en) by Hinoman LTD (https://www.hinoman.com/) with the trade name Mankai, which is reported to have an even higher protein content. Studies with this strain suggest that *W. globosa* is also a good source of bioavailable iron (Yaskolka Meir et al. 2019). As a result, the species has gained attention due its potential as a food product in areas of Europe (van der Spiegel et al. 2013), and is consumed in the US as a powdered supplement.

*W. globosa* and other duckweed species also have other potential uses in sustainable agriculture and phytoremediation. Duckweed’s ability to efficiently absorb nitrogen and phosphorus makes it suitable as a form of soil amendment (Kreider et al. 2019). Research has shown that dried duckweed fertilizer is comparable to inorganic fertilizers while being more effective at reducing nitrogen loss to the environment without sacrificing crop yield (Fernandez Pulido et al. 2024). The nutritional content and high biomass productivity of *W. globosa* also make it an excellent animal feed alternative (Chantirati et al. 2010; Said et al. 2022). In addition, duckweed species also have significant global potential for use in phytoremediation of both heavy metals (e.g., Xie et al. 2013) and eutrophic nutrients (Zhou et al. 2023).

The potential impacts of climate change on the growth and yield of *W. globosa* in its native range are unclear, and at the same time its nutritional profile suggests it could be used as an important food source in these areas. Air temperatures are expected to increase by about 1.5 to 2°C during *W. globosa*’s reproductive months in Thailand, Myanmar (Burma) and Laos in the next 40 years according to the high emissions scenario (SSP5-8.5 scenario; IPCC 2023). Water temperatures on the surface of the small bodies of water where *W. globosa* grows are closely influenced by air temperature (Jacobs et al. 2008). We also know that some crops may be negatively affected by rising mean temperatures (Knox et al. 2012; Bandara and Cai. 2014). In addition, changes to temperature variance and timing are also predicted to generate an increased frequency of heatwave events (e.g., Meehl and Tebaldi 2004). Heatwaves and associated droughts can significantly impact crop yield by decreasing seed size and number, especially when heat events occur during reproductive stages (Hatfield et al. 2011; Cohen et al. 2021).

Thermal performance curves fit to growth rates have been used to predict potential climate change impacts for various crop species (e.g. Agnolucci et al. 2020). Moreover, data on the temperature dependence of growth rates of *W. globosa* would be useful information for commercial production indoors. However, while it is known that temperature is an important driver in duckweed growth rates, *Wolffia sp.* are the among the least studied genera in this group (Pasos-Panqueva et al. 2024). We found no thermal performance curves in literature searches across several databases, including Thai databases, and other reports suggest that there is very limited information about how possible temperature increases and fluctuations will impact the growth rates of *W. globosa* (European Food Safety Authority 2021). There are published estimates of growth rates measured at single constant temperatures between 25 to 28 °C (e.g., Sree et al. 2015; Romano and Aronne 2021); however, since thermal performance is nonlinear, point estimates of growth rates cannot be used to determine growth rates at higher temperatures.

Our objective is to measure the population growth rates of *W. globosa* to temperature in order to understand how predicted increases in Southeast Asia will impact the future for this species as a food crop. Studies on related species such as *Wolffia punctata* suggest that the optimal temperature for this species may be quite high (∼30°C). However, the thermal performance curve of *W. globosa* may have a similar asymmetrical shape as these species, with a sharp drop from the optimal temperature to the upper critical temperature (Docauer 1983). This nonlinear response to temperature will decrease performance when variance is present (Slein et al. 2023), and very sharp asymmetry at higher temperatures could lead to catastrophic impacts of heatwaves. Therefore, we will also examine growth rates under natural fluctuations, and conditions that mimic recent heatwaves in Thailand.

## Materials and Methods

To assess the impacts of climate warming on *Wolffia globosa* performance, we measured the relative growth rate of plants exposed to a range of constant temperatures and fit a thermal performance curve that could be used to identify optimal and suboptimal growing conditions. We compared outcomes between current and future temperature scenarios in Thailand to assess potential impacts of climate change on outdoor production. We also obtained performance under conditions of natural fluctuations, similar to a recent heatwave, to assess the impact of future temperature variance on growth.

### Strain origin and maintenance

Strains of *W. globosa* (accession number 8692) originally collected in Kyushu, Fukuoka, Chikugo, Japan, and curated and stored at Rutgers Duckweed Stock Cooperative (RDSC) were acquired in May 2022 and maintained aseptically. Plants were maintained in 20°C Memmert incubators (IPP-500) and cultivated on a modified Hoagland’s medium, a macronutrient solution commonly used for culturing duckweed species (e.g., Fang et al. 2020). A 16:8 light-dark photoperiod was applied using LED lights (Kema Keur 0J-304 250V-3A 3A/125VAC E354212) to provide daylight intensity between 38.6-44.3 µmol m−2 s−1, which is a moderate level for the growth of duckweeds under controlled conditions (Pasos-Panqueva et al. 2024). About every 20 days, we created a new source population using new growth media to ensure that plants selected for experiments maintained active metabolic rates.

### Temperature treatments

We used a range of constant temperatures (14.0°C, 20.0°C, 25.0°C, 26.5°C, 28.5°C, 31.5°C, 37°C) to construct a thermal performance curve. The lowest temperature tested, 14°C, was selected because it corresponds to the threshold at which similar species typically enter dormancy, producing inactive structures called turions (Romano and Aronne 2021). We selected intermediate temperatures to allow comparison to both previously published work (e.g., Ziegler et al. 2015), and to match current and predicted mean temperatures during the growth season in Thailand, as well as mean temperatures during a recent heatwave (see below for description). The highest temperature of 37°C was chosen based on the upper critical threshold for two other tropical species, *Wolffia punctata* and *Wolffia columbiana* (Docauer 1983).

To assess how predicted temperature increases will impact *W. globosa* production, we focused on Thailand’s current and future climate, as this country has a longstanding tradition of *W. globosa* production (Siriwat et al. 2023). We used monthly historical (1991 - 2020) and predicted (2040 - 2059) average temperatures from the World Bank Climate Change Knowledge Portal (2024), as data at finer temporal scales is not available. Within this portal, predicted temperatures are based on the 2023 Coupled Model Intercomparison Projects (CMIPs) from the Intergovernmental Panel on Climate Change (IPCC 2023).

We identified the current average temperature during the growing season (November to July), the period when *W. globosa* is predominantly grown (European Food Safety Authority Panel on Nutrition, Novel Foods, and Food Allergens 2011). This value represents the mean of the monthly 50^th^ percentile (median) temperatures for that period. Next, we applied the SSP5-8.5 climate change scenario, which is a business-as-usual emissions prediction without additional mitigation efforts (IPCC 2023), to estimate the average temperature increase in Southeast Asia. By averaging the predicted monthly 50th percentile (median) temperatures temperature increases for the growing season months, we determined that mean temperatures are expected to rise by approximately 2°C over the next 20–40 years, raising Thailand’s average growing season temperature from 26.5°C to 28.5°C (The World Bank Group 2024).

To assess how warmer temperatures with natural variation might impact *W. globosa* performance, we also created two fluctuating treatments based on the following reference scenarios: 1) mean current temperature (26.5°C); and 2) mean temperature (31.5°C) during a heatwave in Thailand from April 22 to May 23, 2024 (Visual Crossing Corporation 2024; see Supplementary Information 1). For the mean current temperature, we obtained the average minimum and maximum monthly temperatures for the November-July period. For the heat wave scenario, we used the average hourly minimum and maximum temperatures, along with the overall hourly average temperature. We adjusted the minimum and maximum temperatures to ensure that the average temperatures in the fluctuating treatments matched those of the constant temperature treatments, while preserving a consistent standard deviation of 3.2°C to 3.3°C across the two fluctuating treatments (Table 1).

**Table 1.** Temperature parameters, based on historical and predicted temperatures in Thailand, used to create fluctuating thermal sequences for each treatment: season average and heatwave.

| **Temperature** | **Season Average** | **Heatwave** |
| --- | --- | --- |
| Minimum | 21.2 °C (22 °C) | 28.3 °C (27.1 °C) |
| Mean | 26.5 °C | 31.5 °C |
| Maximum | 31.9 °C (31 °C) | 36.1 °C (36 °C) |
| Standard deviation* | 3.3 °C | 3.2 °C |

\* Standard deviation values were calculated based on thermal sequences generated using the adjusted minimum and maximum temperatures (listed in parentheses).

We used the adjusted maximum and minimum temperatures to create fluctuating temperature treatments using a sinusoidal function, a commonly employed method for simulating temperature cycles (National Institute of Water and Atmospheric Research 2023), using the following equation: *T(hour) = min + (max - min) * (0.5 + 0.5 sin (2π (hour) / 24)),* where T(hour) is the temperature at a given hour, hour is the index representing each hourly time step, and min and max are the minimum and maximum temperatures in each scenario. The sinusoidal function generates thermal sequences with hourly temperatures by interpolating between the minimum and maximum using a sine wave, ensuring smooth transitions over a 24-hour period to mimic natural daily temperature fluctuations. For each treatment, we repeated this pattern daily for a total of 5 days (Figure 1).

**Figure 1.**
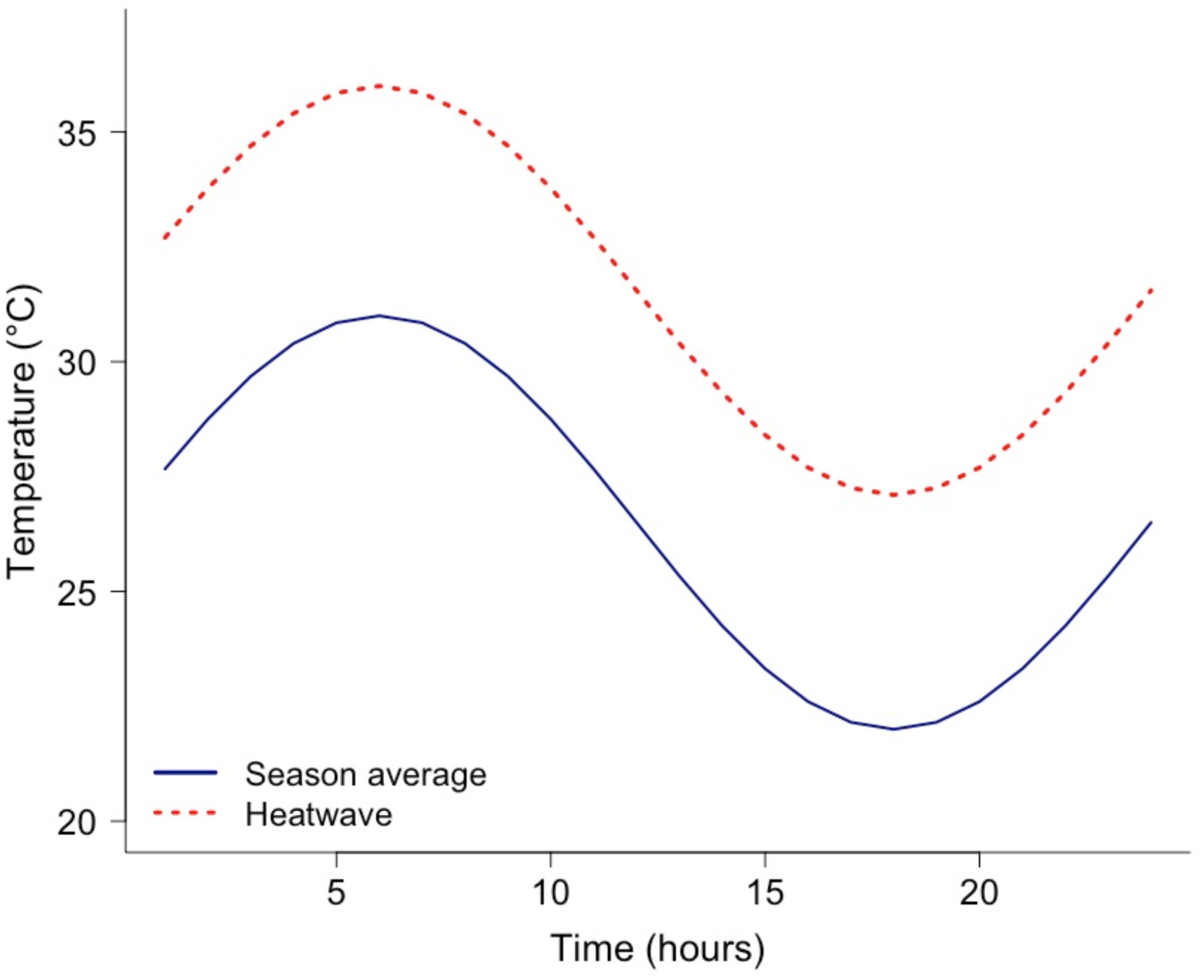
Daily fluctuation for variable temperature treatments. The season average treatment reflects the current growing season temperatures in Thailand, and heatwave treatment mean and variance are based on an actual heat wave event that occurred from April-May 2024.

### Experimental methods for temperature treatments

The experiments were conducted over a five-day (120-hour) period, which was sufficient to observe a change in the number of fronds. Temperature treatments were conducted in the previously described incubators, using the proprietary software Celsius (version 10.1.4) to implement the desired sequence of temperatures. Each replicate consisted of twelve pairs of fronds randomly selected from a source population of *W. globosa* and placed into a 24 well-plate, with each well containing about 1000 µL of Hoagland’s solution. Replicates of each temperature treatment were randomly assigned to individual incubators. For each temperature treatment, a minimum of five replicates were completed.

At the start and conclusion of the 5-day experiment, a Zeiss Stemi 2000-C stereomicroscope was used to examine frond health and perform count fronds for each replicate. Fronds were classified as alive if they exhibited less than 90% loss of green pigmentation (Thomson and Dennis 2013); otherwise, they were considered dead. Fronds of all sizes were included, and both live and dead fronds were counted.

### Data analysis

We fit three thermal performance curves to the relative growth rate across all constant temperature treatments. We used a modified Ratkowsky model as: *(a(T-Tmin))^2^(1-e(b(T-Tmax)))* where *T* is the temperature, *Tmin* is the low temperature where rate of performance becomes 0, *Tmax* is the high temperature where rate of performance becomes negative, and and *b* are rate parameters (Ratkowsky et al. 1983). We also fit a Brière2 curve (Brière et al. 1999) given as: *aT(T - Tmin)(Tmax-T)#x005E;(1/b),* and a simple quadratic urve We fit these curves using the R package *nls-multstart* (Padfield and Matheson 2023) where minimum and maximum values for initial coefficients were Tmin=(-15, 5), Tmax=(14, 37), a=(-0.5, 1.5), and b=(-1, 1.1), for both the Ratkowsky and Biere2 functions, and a=(-2,2), b=(-2,2), and c=(-1,1) for the quadratic.

We used generalized linear models with a Poisson distribution to analyze the impact of temperature treatments on the number of live and dead fronds, where replicates were randomly subsampled to ensure an equal number in each temperature treatment. All analyses were performed using R Statistical Software (v3.6.3; R Core Team 2020), using the package (Russell 2021) for pairwise treatment comparison.

## Results

The best fit thermal performance curve possesses a clear asymmetrical shape, with a drastic decline in growth rate towards the higher temperatures, with a suggested optimal temperature ∼ 32°C (Fig 2). The best fit curve for the relative growth rate was the modified Ratkowsky function as indicated by AIC (Ratkowsky: -102.4, Brière2: -101.8 and quadratic: -85.3) and the residual sum of squares (Ratkowsky: 0.31, Brière2: 0.31 and quadratic: 0.45). Although the difference in the fit of the Ratkowsky and Brière2 curves is negligible.

**Figure 2.**
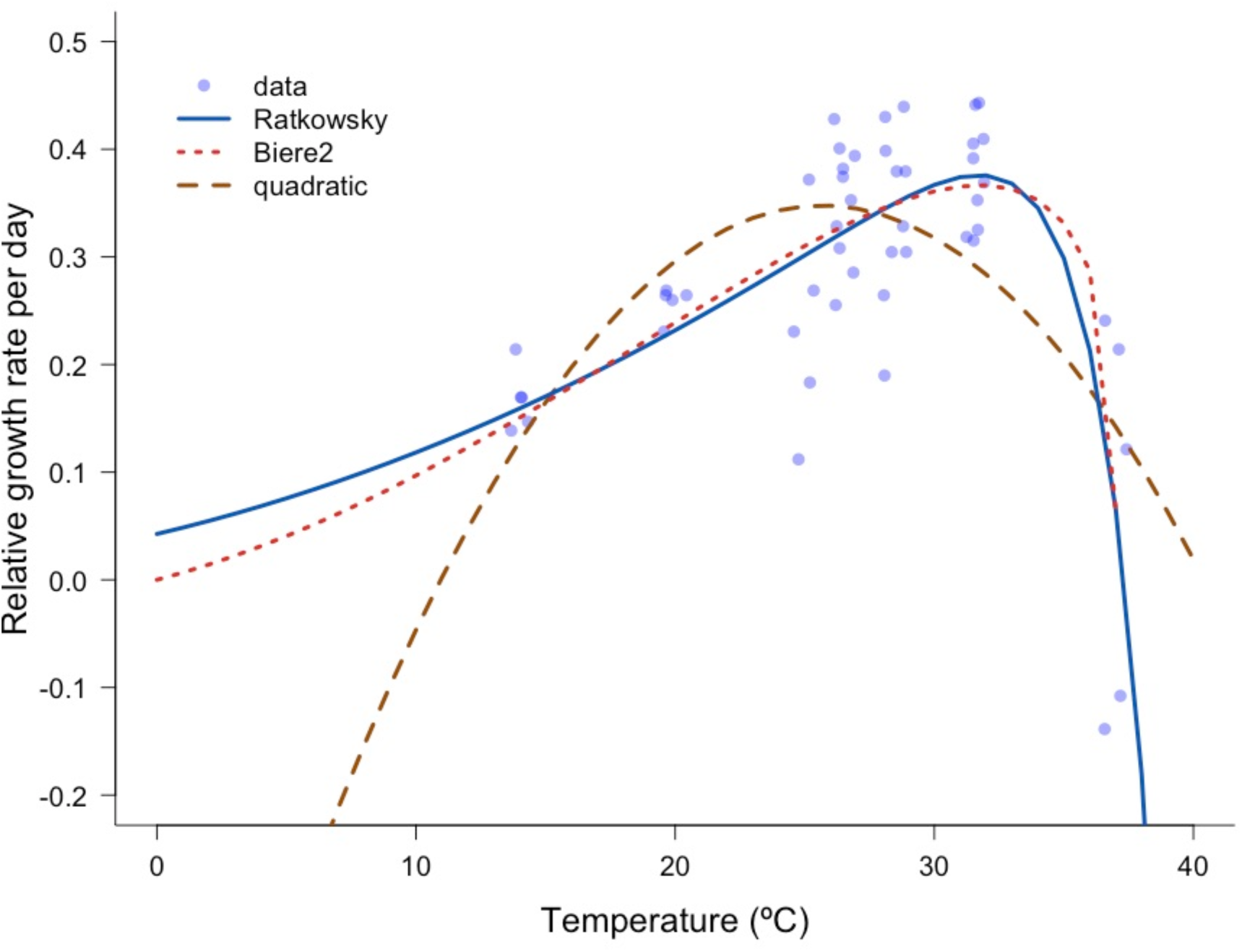
Modified Ratkowsky, Brière2 and quadratic functions fit to the relative growth rate of *W. globosa* measured at different constant temperatures. The modified Ratkowsky curve has the best fit with parameters a= 0.014, b=0.43, Tmin=-15°C and Tmax=37.3°C. A small amount of jitter has been added to the data to improve clarity.

Temperature affected the number of live fronds after 5 days (Fig 3a). The number of fronds in the lowest (14°C) and highest (37°C) constant temperature treatments was significantly smaller than all other temperature treatments (p<0.0001 for all pairwise comparisons), but was not significantly different between these two treatments (z=-1.893, p=0.058). However, there was no significant difference between the number of fronds in treatments with temperatures at the current mean during the growing season in Thailand (26.5°C) and the projected mean in 40 years (28.5°C) (z ratio = 0.150, p=1.00).

**Figure 3.**
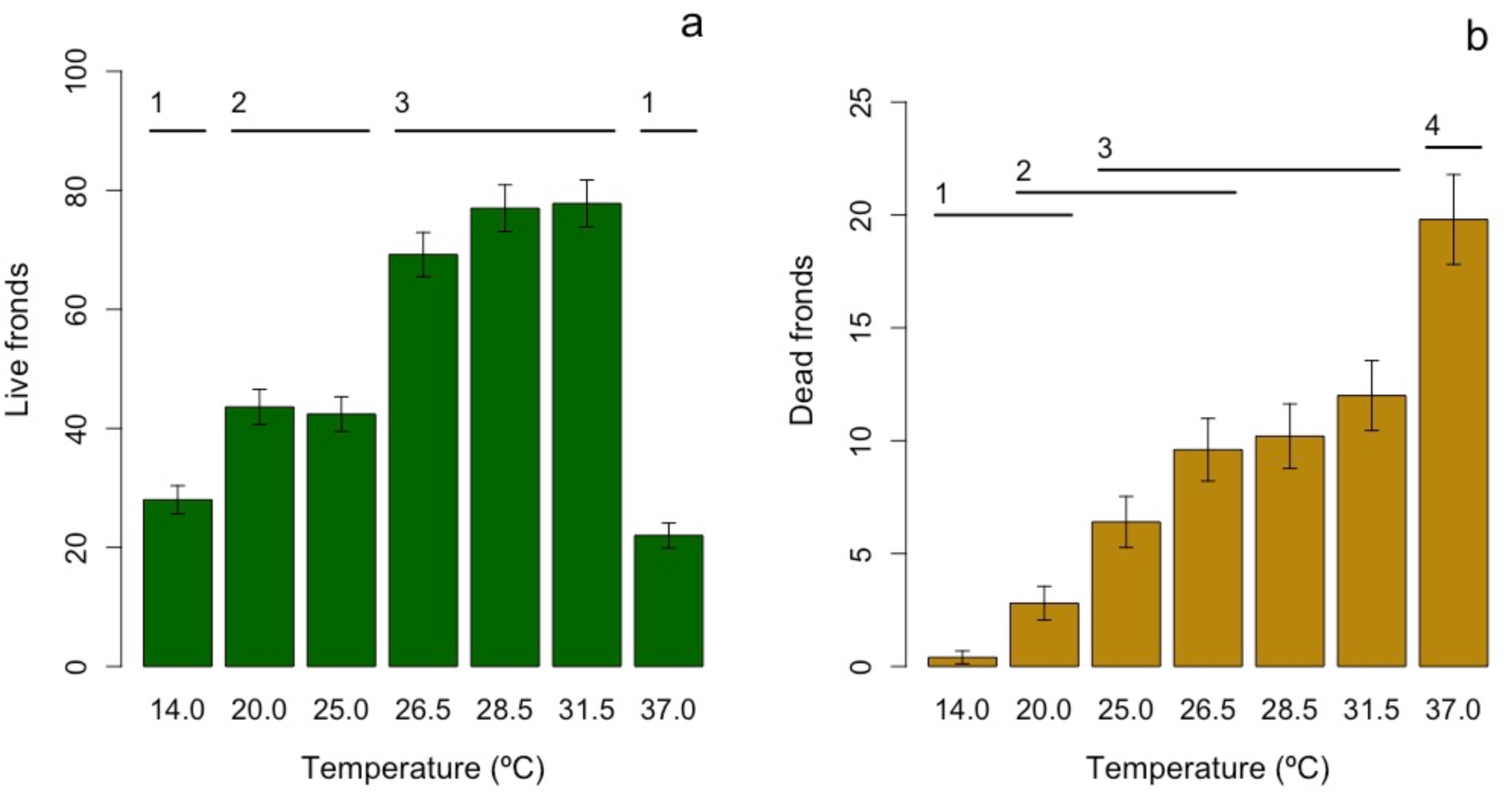
Mean and standard error calculated using a Poisson distribution of the (a) number of live fronds or (b) number of dead fronds of *W. globosa,* where there are 5 replicates at each of the different constant temperatures. Horizontal bars and numbers indicate temperature treatments that are not significantly different.

There was a significant impact of temperature on the number of dead fronds. Therefore, although the number of live fronds did not differ in the coldest and hottest temperature treatments, analysis of the number of dead fronds clearly indicated that there were different mechanisms at work: reduced reproductive rate at cold temperatures and increased death rate at high temperatures. The largest number of dead fronds was found in the hottest treatment (37°C, all pairwise comparisons p<0.05) and the smallest number was in the two coldest treatments (14°C and 20°C; these two treatments are not significantly different; z.ratio=-2.574, p= 0.1339) (Fig 3b). Again there was no difference in the number of dead fronds at the current mean temperatures during the growing season in Thailand (26.5°C) and the projected mean in 20-40 years (28.5°C) (z.ratio=-1.388, p=0.8083).

We note, however, that constant temperature treatments may not reflect the impact of naturally fluctuating temperature regimes. Comparisons between the number of live fronds in either constant or fluctuating temperature treatments at either 26.5°C or 31.5°C indicated that there was a significant interaction (p<0.0001) between temperature and constant vs fluctuating regimes (Fig 4). The number of live fronds was lower in the 31.5°C temperature treatment when fluctuations were present, but higher under constant conditions.

**Figure 4.**
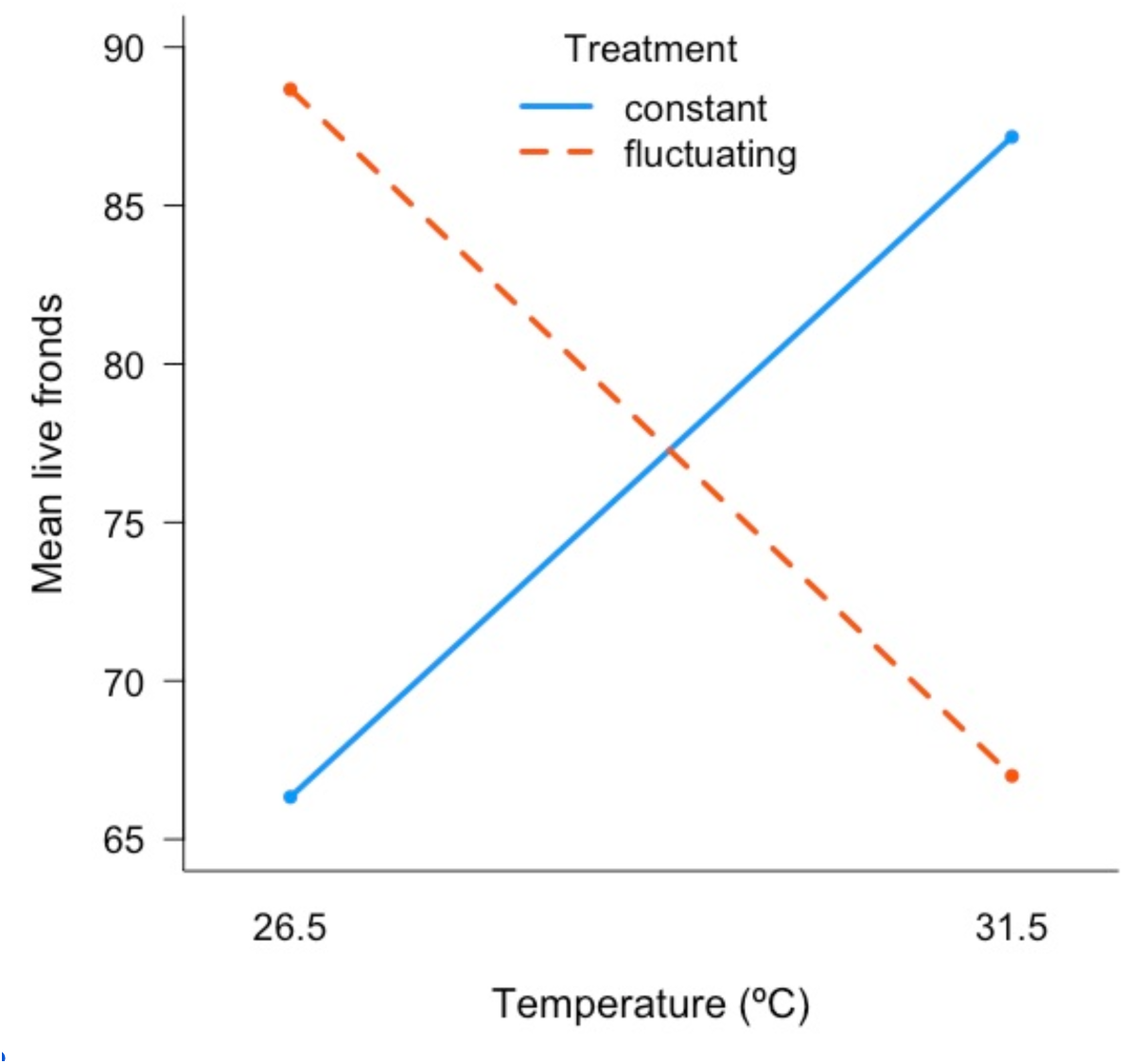
Interaction plot of the mean count of live fronds of *W. globosa* where there are 6 replicates for each treatment. Temperatures are either constant (blue lines and points), or fluctuating (dashed red lines and points).

## Discussion

The relative growth rates of *Wolffia globosa* may have an optimum at higher temperatures than expected. However, the best fit thermal performance curve is quite nonlinear and asymmetric, suggesting substantial decreases in growth at temperatures higher than the optimum. At constant temperatures reflective of maximums during recent heatwaves, relative growth rate was reduced because of higher frond death rates. When temperatures are allowed to fluctuate, the asymmetric shape of the curve caused greater reductions in growth near the optimum temperature, so that under natural variation higher growth rates are expected at cooler temperatures. Therefore, it is likely that future climate change impacts, such as increased frequency of heatwaves and increased mean water temperatures, may have a negative impact on outdoor cultivation of the species.

The highest measured relative growth rates were at 31.5°C Growth rates range from a minimum of 0.17± 0.033 per day (at 14°C) to a maximum 0.38± 0.023 (at 31.5°C) per day, and are similar to previously published rates for *W. globosa* Romano and Aronne (2021) indicate that the relative growth rate of *W. globosa* ranges from 0.155 to 0.559 per day. Sree et al. (2015) reported a relative growth rate of 0.155 ± 0.040 per day, while Ziegler et al. (2015) reports a rapid relative growth rate of 0.328 ± 0.022 per day for *W. globosa* at 25°C. We note that growth rates are highly dependent on the particular clone, nutrients and light levels (e.g., Sree et al. 2015), which vary amongst these studies.

We provide the first thermal performance curve for *W. globosa.* The best fit curves have a typical asymmetric shape, rather than symmetric one, with slow decline in relative growth rates as the temperature becomes colder, but an abrupt decrease in performance at warmer temperatures (Fig 2). Both fitted asymmetric curves suggest that, at constant temperatures, the maximum relative growth rate occurs at ∼32°C, while growth decreases below 26.52°C and above 32°C. More measurements at temperatures greater than 31.5°C would increase our confidence in this estimated optimum. The thermal performance for *W. globosa* is quite similar to that of some other species of duckweed, such as *Wolffia punctata* and *Wolffia columbiana* (Docauer 1983). These species also have an asymmetric response to temperature with an optimum near 30°C and an abrupt decrease with an upper critical threshold near 37°C (Docauer 1983).

An analysis of the number of live and dead fronds reveals that there are different causes for the decreased relative growth rates at low and high temperatures. While there is no difference in the number of live fronds at the highest and lowest temperatures tested, there were significantly more dead fronds at the highest temperature. In fact, at the coldest temperature there were nearly zero dead fronds. Therefore the reduced relative growth rates at cold temperatures are primarily due to reduced reproduction, while the lower or negative growth rates at hot temperatures are caused by an increased death rate. While higher temperatures are known to reduce duckweed lifespan (Paiha 2021), the short length of our experiments and the large numbers of dead fronds point to temperature-dependent mortality rather than accelerated senescence as the cause of the reduced growth rates at the highest temperatures.

This information regarding the temperature dependence of the relative growth rate of *W. globosa* has implications for both indoor cultivation and also its spread in non-native regions. Recently there has been increased interest in commercial production of the species, and for its use in space travel (Romano and Aronne 2021). For example, In 2021, the Israeli company (https://www.greenonyx.ag) opened a novel food dossier for both *W. globosa and W. arrhizia* The consumption of *W. globosa* was approved by the European Food Safety Authority. Recommendations regarding cultivation typically suggest a broad range of conditions from 20°C to 30°C as optimal (Kuar 2024), but our studies indicate that maximum growth will occur in a narrower and warmer range from 26.5 to 32°C. Nevertheless, it is clear that the species has positive growth rates at the coolest temperature measured (14°C), and therefore, it is likely to continue its spread through regions of Europe and North America with moderate temperatures (e.g., UK: Lansdown et al. 2022, Germany: Frank et al. 2020 and France: Lecron et al 2021), where like other invasive duckweeds, it may have negative impacts (e.g., Ceschin et al. 2020).

Our thermal performance curve with constant temperatures could suggest that there will be no significant near term impacts of climate change on the growth of *W. globosa* in its current areas of outdoor cultivation. We find no significant difference in the growth of *W. globosa* at the current mean air temperature during the growing season in Thailand (26.5°C) and the projected mean in 40 years (28.5°C) of the 2023 Coupled Model Intercomparison Projects (CMIPs) from the Intergovernmental Panel on Climate Change (IPCC, 2023). In comparison, the predicted increase in temperature in Laos and Myanmar, from 23.5°C to 25°C (The World Bank Group 2024), will bring mean temperatures closer to *W. globosa*’s optimum; therefore possibly increasing its growth rate. Over a longer timeframe, under the high-emission SSP5-8.5 scenario for Southeast Asia, projections suggest that average July surface air temperatures in the region could rise to 27.62°C by 2050 and 29.13°C by 2100 (Sentian et al. 2022), but this is still within the range of optimal performance.

However, when there is a nonlinear thermal response with an abrupt decrease, even small variance in temperature conditions can decrease performance (Slein et al. 2023). We found a significant interaction when we compared constant and fluctuating temperature regimes. When temperatures were allowed to fluctuate, the relative growth rate was higher at 26.5°C rather than 31.5°C. Therefore it is possible that when a realistic level of temperature fluctuation is considered, climate change may result in growth decreases. This conclusion is reinforced when we consider that a greater frequency of heatwaves is anticipated under future conditions (Meehl and Tebaldi 2004). Thailand’s temperature fluctuations and extremes on a local scale are almost linearly related to large-scale or global mean temperature changes (Amnuaylojaroen and Parasin 2022). The mean and degree of variance of the 31.5°C fluctuating treatment was based on actual air temperatures experienced during the 2024 heatwave in Thailand. This temperature treatment had mean conditions not very different from both the mean water temperatures in fish ponds in May (Sriyasak et al. 2013), and the projected summer mean air temperature for Thailand in 2080 (30.63°C, SSP5-8.5; The World Bank 2024). Therefore, if current emission trends persist, temperature conditions similar to our heatwave treatment will increase, and growth rates may decline.

At the water surface where *W. globosa* grows, we might expect mean temperatures to be very closely linked to mean air temperatures (Jacobs et al. 2008). However, the variance of air temperatures is usually larger than water temperatures because the high specific heat capacity of water dampens fluctuations. In addition, in deeper ponds where stratification is possible, water temperatures may be cooler than air temperatures. For example, a study of 18 fish ponds with depths of 0.8 - 2.0m in northern Thailand reports that during the dry season in January, mean air temperature was warmer (28.3°C ± 4.11°C) and had a much wider range (6.5°C-35.83°C) than water temperatures (mean of 26.3°C ± 0.57°C with range from 25.5°C to 27.1°C) (Sriyasak et al. 2013). The rainy season decreases stratification, and in May, mean water temperatures were warmer than mean air temperatures (mean air temperature: 28.14°C ± 4.03°C; mean water temperature: 30.44°C ± 0.80°C). Therefore, water temperatures always had lower variance, and the constant temperatures used to construct the thermal performance curve may reasonably reflect this lack of fluctuation. However, the mean water temperatures reported during the rainy season are right at the optimum for *W. globosa.* At these temperatures, even a small amount of variance could lead to abrupt decreases in growth rates. Moreover, in the Lower Songkhram River regions, where *W. globosa* used for food (Norcum et al. 2025), water temperatures may rise beyond the optimum (e.g., 35°C) during the growing season (Satrawaha et al. 2009). Future research on the thermal performance of *W. globosa* should include information on seasonal differences in both the mean and variance of water temperatures at the surface of these small water bodies.

Other impacts of climate change, not considered here, may also decrease outdoor production. Drought is associated with climate change and heatwaves (Khadka et al. 2024) and may impact the occurrence of ponds and small waterbodies that are required for cultivation. Intraspecies competition with other surface-covering plant species, such as algae, cyanobacteria, and other duckweeds is common (Coughlan et al. 2022, Ziegler et al. 2023), and may increase with climate change. For example, Gillies et al. (2024) find that high-temperature stress could increase sensitivity to competition in Lemna duckweed species.

Predictions for other crop species in this region, such as rice and corn, similarly suggest that near term increases in mean temperatures may benefit growth rates, but that extreme conditions would decrease yield (e.g., Amnuaylojaroen and Parasin 2022, Waqas et al. 2025). While *W. globosa* is likely to be negatively impacted by larger increases in mean temperatures and associated heatwaves, we should note that its very high growth rate at average high temperatures will enable quick recovery from extreme conditions. Depending on the timing of the event, crops with comparable protein, such as peas or soy may have a reduction or complete loss of harvest for the season (e.g., Siebers et al. 2015). In contrast, as long as a small proportion of the fronds survive, *W. globosa* has the potential for ample future harvests in a few weeks.

## Conclusions

There are a limited number of studies evaluating thermal performance curves of plants (Wooliver et al. 2022), even though such data may provide crucial information for climate change adaptation. We provide the first thermal performance curve of the relative growth rates of *W. globosa*, and use this information to predict responses to climate change in Southeast Asia where it is currently cultivated. While performance under constant temperature conditions suggests little impact under climate change, we find decreased growth rates when temperatures fluctuate, and note that the highly asymmetric thermal performance curve can lead to sudden catastrophic impacts at high temperatures. Nevertheless, the fast growth rate of the species suggests fast recovery from heat events is possible. This potential for resilience and the excellent nutritional profile of the species suggest it could play an important role in climate change adaptation.

## Data Availability

Relative growth rates for *W. globos*a and R code for data analysis will be available after publication at https://github.com/Cuddington-Lab

## Supplementary Information 1

**Table.** Daily temperatures from a heatwave in Thailand. Data obtained from Visual Crossing Corporation (2024).

| Date | Minimum (°C) | Maximum (°C) | Mean (°C) |
| --- | --- | --- | --- |
| 4/22/2024 | 30 | 37.5 | 33.3 |
| 4/23/2024 | 30.2 | 37.5 | 33 |
| 4/24/2024 | 30 | 37.7 | 32.9 |
| 4/25/2024 | 29.7 | 38.7 | 33.1 |
| 4/26/2024 | 29.2 | 37.5 | 32.9 |
| 4/27/2024 | 30 | 38.2 | 33.3 |
| 4/28/2024 | 30.4 | 37.7 | 33.2 |
| 4/29/2024 | 30.4 | 38.5 | 33.4 |
| 4/30/2024 | 30.6 | 40 | 33.8 |
| 5/1/2024 | 31 | 38.2 | 33.9 |
| 5/2/2024 | 31 | 38 | 33.5 |
| 5/3/2024 | 30 | 38.5 | 33.5 |
| 5/4/2024 | 30.1 | 36.7 | 32.7 |
| 5/5/2024 | 30 | 39.1 | 33.4 |
| 5/6/2024 | 29.8 | 37.4 | 32.8 |
| 5/7/2024 | 24.7 | 31.4 | 27.2 |
| 5/8/2024 | 25.1 | 35.9 | 30.1 |
| 5/9/2024 | 28.4 | 35.6 | 31.1 |
| 5/10/2024 | 26.9 | 35.7 | 30.8 |
| 5/11/2024 | 26.7 | 33.4 | 29.8 |
| 5/12/2024 | 26.9 | 36.6 | 30.3 |
| 5/13/2024 | 25 | 34.6 | 29.3 |
| 5/14/2024 | 25.3 | 35.8 | 30.2 |
| 5/15/2024 | 28.2 | 36.7 | 32.1 |
| 5/16/2024 | 28 | 34.1 | 30.1 |
| 5/17/2024 | 28.8 | 35 | 31.3 |
| 5/18/2024 | 28.2 | 33.7 | 30.9 |
| 5/19/2024 | 25.9 | 34.3 | 30.3 |
| 5/20/2024 | 26.2 | 35.7 | 30.4 |
| 5/21/2024 | 26.4 | 33.1 | 29.4 |
| 5/22/2024 | 26.6 | 34 | 29.6 |
| 5/23/2024 | 26.2 | 30.5 | 28 |

